# Deep brain cancer metastasis detection with wide-volume Raman spectroscopy through a single tapered fiber

**DOI:** 10.1101/2022.06.24.497456

**Authors:** Filippo Pisano, Mariam Masmudi-Martín, Maria Samuela Andriani, Elena Cid, Marco Pisanello, Antonio Balena, Liam Collard, Francesco Tantussi, Marco Grande, Leonardo Sileo, Francesco Gentile, Francesco De Angelis, Massimo De Vittorio, Liset Menendez de La Prida, Manuel Valiente, Ferruccio Pisanello

## Abstract

We propose a low-invasive method to enable implantable, large volume Raman spectroscopy in arbitrarily deep brain regions of the mouse brain. Using a single tapered fiber as thin as 1 μm at the tip, we identified diagnostic markers of brain metastasis - the most frequent brain tumor in human adults - with overall accuracy ≥ 90%. We view our approach as a promising complement to the existing palette of tools for optical interrogation of the brain.

## Main

Over the last decade, developments in neuro-technology have been aimed at controlling and monitoring neural activity over wide cellular populations. Electrophysiological recordings can now sense and sort signals from thousands of neurons simultaneously (*1*), while all-optical methods based on genetically encoded actuators and reporters have introduced cell-type specificity in functional connectivity studies as well as time-resolved imaging of neurotransmitters transients (*2*–*4*).

However, to complement this picture the field still lacks *label-free* approaches to detect bio-molecular changes from large volumes in deep brain regions. The progression of several types of brain pathologies is indeed known to involve slow but permanent bio-chemical alterations, including Alzheimer’s disease (with the emerging of beta amyloid plaques) and brain metastasis. To achieve this sensing capability, one would naturally turn to vibrational spectroscopy (e.g., Raman spectroscopy), a suite of techniques which has found countless applications in life science (*5*). Nonetheless, the use of implantable Raman in neuroscience is still in its infancy and presently requires exogenous reporters (*6*). To exploit its full potential, the field needs appropriate technologies with minimal invasiveness and large collection volumes suitable for small animal models, such as the mouse. While advanced Raman endoscopes have been developed to provide intraoperative guidance during tumor resections (*7*–*9*), these probes are too bulky to work as neural implants in laboratory species as they require multiple waveguides to increase the recording volume and to disjoin excitation and collection signals, suppressing the probe’s Raman response.

To tackle this challenge, we view tapered optical fibers (TFs) as a promising implantable optical system. Owing to the mode-division de-multiplexing properties of a narrowing diameter section (*10*), TFs can exchange photons with the surrounding tissue over implant depths greater than 1 mm and have already been employed for large-volume optical neural interfaces to control and monitor neural activity in freely moving mice (*11, 12*). The narrowing diameter of these implants (from 230 μm at its widest to 1 μm at the tip) reduces tissue reaction with respect to standard fiber optics (*13*) and establishes photonic properties that support a wide palette of optical detection strategies (*14, 15*) from the visible to the near infrared range.

In this work, we devise a method to probe Raman signals at depth from cortical to subcortical structures of the mouse brain, getting label-free access to biomolecular markers on a wide tissue volume around and along the implant to detect the presence of brain metastasis (Fig. 1a).

**Figure 1.**
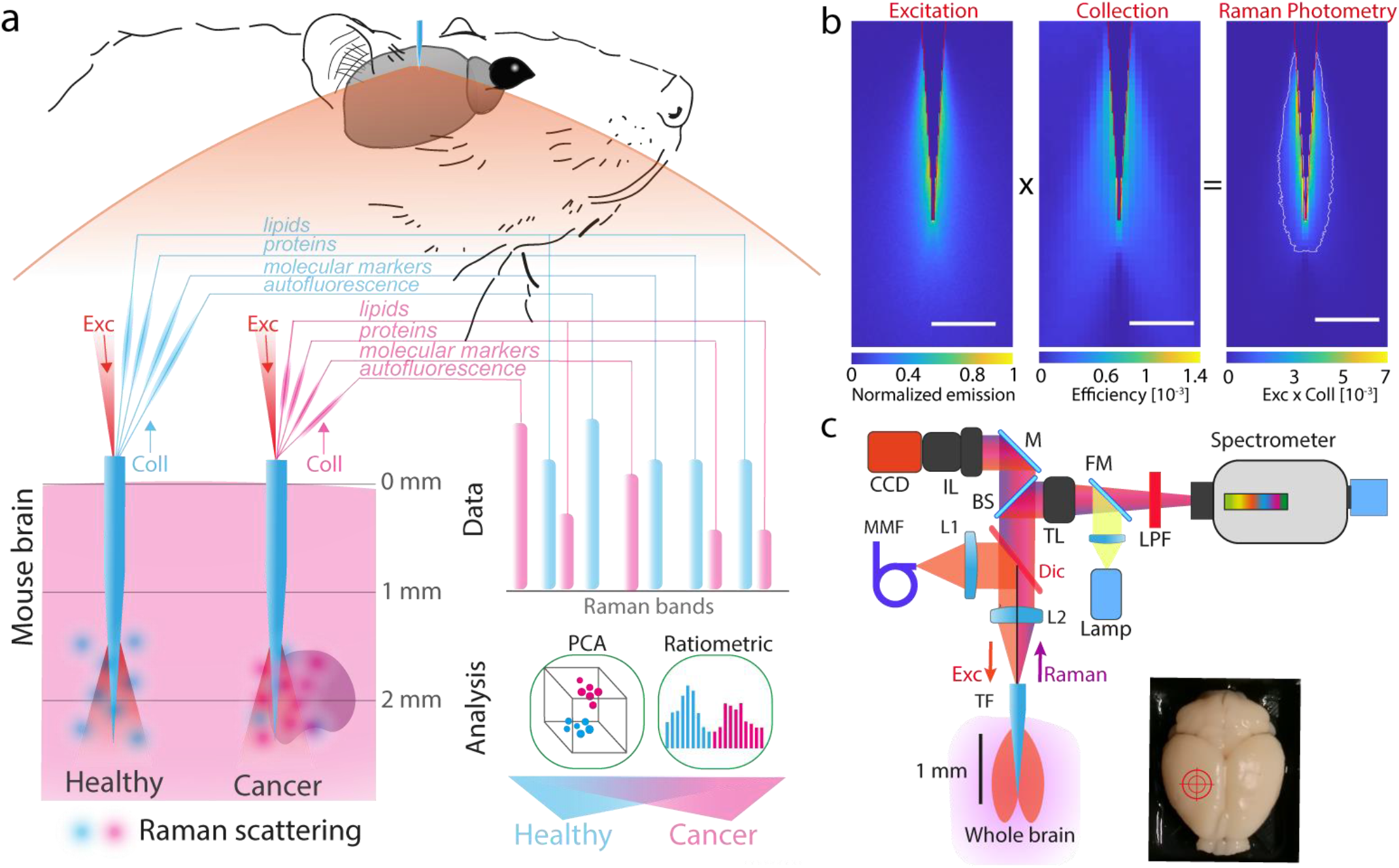
**a**, Concept of the approach: minimally-invasive implantable tapered fibers (TF) collect information on multiple biomarkers from the Raman response of the surrounding volume of tissue in deep brain regions, lead to the detection of metastasis via Principal Component Analysis (PCA) or Ratiometric analysis. **b**, (left) Numerical simulations of the excitation volume at 785 nm, the illumination intensity is scaled to 1; (center) a representative collection volume in the first NIR window (calculated for scattering at 920 nm), the color bar shows the collection efficiency; (right) the Raman Photometry volume calculated from the emission and collection ones (see Methods); all scale bar 500 μm. **c**, Optical system to excite and collect Raman signals through the implantable TF. Full details in the methods section (MMF, multimode fiber; L1, L2 lenses; TF tapered fiber; Dic, dichroic mirror, BS, beam splitter; IL, imaging lens TL, tube lens, FM, flip mirror, LPF, long pass filter, M mirror, CCD Charge Coupled Device).

To alleviate the impact of the Raman background from the fiber without increasing the size of the implant, we maximized light collection using a single waveguide to deliver the excitation (785 nm). We collected signals in both the fingerprint (FP, 800-1750 cm^-1^) and the high wavenumber (HW, 2800-3100 cm^-1^) spectral ranges targeting the lipid, protein and other molecular bands (for example melanin). This broad spectral range can be harnessed through the tapered fiber because the extension of the optically active region remains unaltered despite the number of guided modes decreasing with the wavelength. This allows to deliver and collect excitation light and Raman scattered photons along the same dorso-ventral extent. The volume of tissue that contributes to the Raman signal is shown in Fig. 1b as a combination between the illuminated volume of tissue, calculated at 785 nm, and the collection volume, calculated for radiation scattered at the center of the spectral collection range (920 nm or 1870 cm^-1^, see Methods)(*16*). This is hereafter referred to as Raman photometry volume. To experimentally detect this signal we devised a custom microscope designed around a TF implant stub to enhance signal-to-background ratio (Fig. 1c, see Methods).

First, we measured the variations in the Raman spectra acquired in different anatomical regions when inserting the probe at increasing depth in the dorso-ventral (DV) direction of naïve mouse brains (Fig. 2a see Methods). As the TF travelled from cortical to sub-cortical brain regions (Fig. 2b), the Raman spectra in the HW range reflected the variations in the local molecular composition (*17, 18*) (Fig. 2c). The presence of cells’ somas in cortical layers is highlighted by strong protein bands (2934 cm^-1^) when recording at shallow depths (from 0 mm to -1 mm in the DV axis); on the other hand, lipid signatures (2845-2875 bands cm^-1^) become more prominent in deeper regions while recording through the *corpus callosum*, the fimbria of the Hippocampus (HP) and myelinated regions around the Thalamus (TL) (e.g., from -1 mm to -2 mm and from -2 mm to -3 mm in the DV axis).

**Figure 2.**
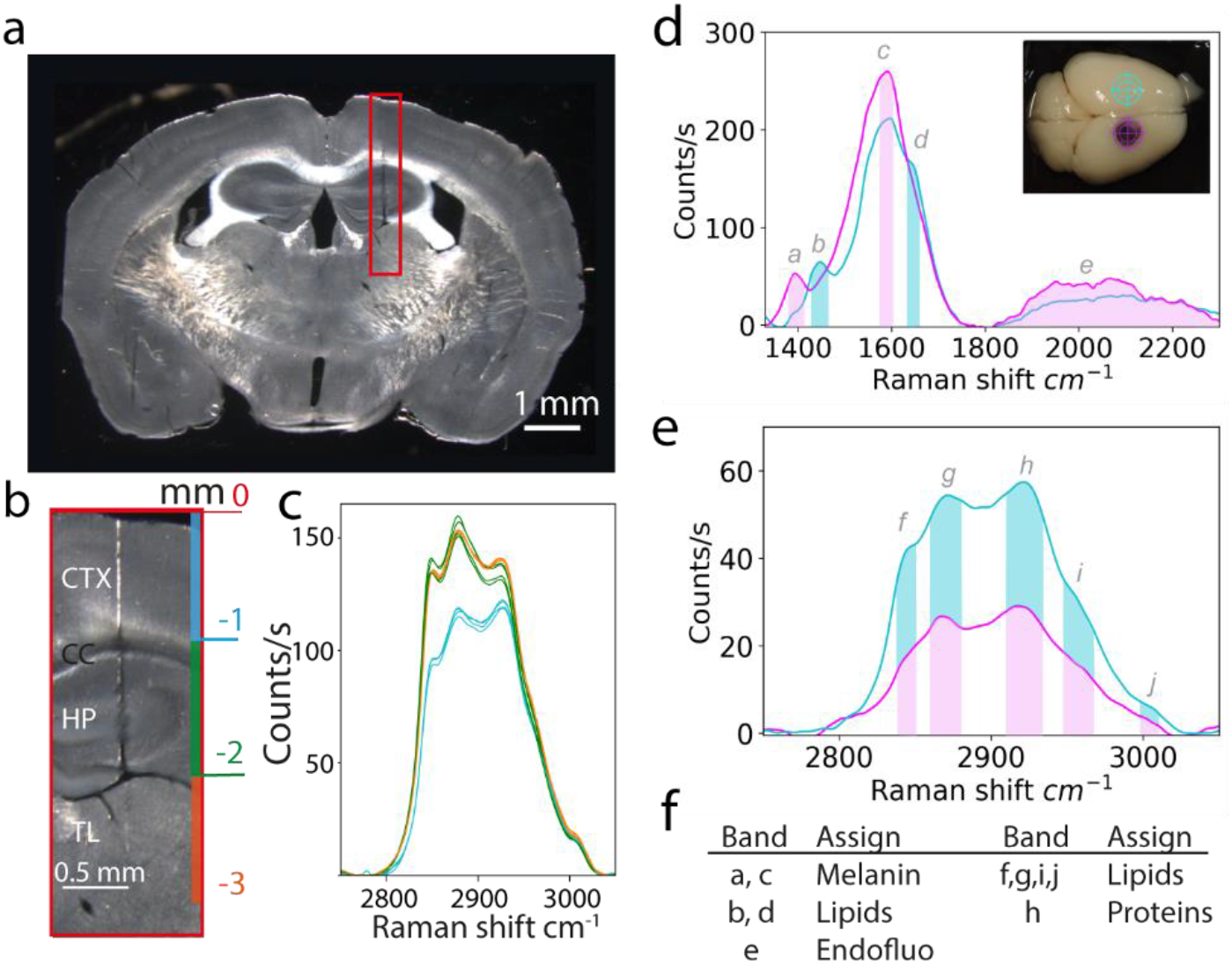
**a**, Wide field image of a brain slice with the TF track **b**, Close up of the fiber track with relevant anatomical regions and collection depths **c**, Raman spectra acquired at progressive depths, colors refers to panel b, **d, e** Representative Raman spectra acquired in a metastasis (magenta) and in a contralateral position (cyan) with the relevant bands highlighted **f**, attribution of the Raman bands

Then, we focused on detecting the presence of B16/F10-BrM melanoma brain metastasis (*19*) based on the spectral features extracted by the TF. We performed a systematic study inserting the probes in four whole brains with melanoma brain metastases and three control brains. For each brain, we recorded the Raman signal at progressively deeper locations at the core of the lesion, at the lesion’s periphery and in the contralateral hemisphere, at a symmetric position on the medio-lateral axis with respect to the lesion’s core. Similar implant positions were used for control brains. We acquired signals in the FP and HW ranges, taking 5 measurements at each position (see Methods). Fig. 2d, e show representative averaged Raman spectra collected in cancer free regions (cyan) and regions with metastatic colonization (magenta) in the FP and HW ranges.

We identified Raman signatures that correlate with brain tissue affected by metastasis including the insurgence of peaks compatible with melanin pigment in cancer cells (bands *a* 1400 cm^-1^ and *c* 1590 cm^-1^, Fig. 2d) (*20*), a diminished signal in the FP lipid bands (bands *b* 1440 cm^-1^, *d* 1650 cm^-1^ Fig. 2d) (*8*), a diminished signal of CH (Carbon-Hydrogen) bands in the HW range (*f, g, h, i, j* 2850-3000 cm^-1^ in Fig. 2e) (*8, 21*) and a sizeable auto-fluorescence contribution across the Raman silent region (band *e* 2000-2500 cm^-1^ in Fig. 2e) (see Fig.2f).

Then, we segmented the spectra using Principal Component Analysis (PCA). PCA was used to reduce the dimensionality of the spectra while preserving much of the information (99%) of the original datasets (*22*). The three most informative components (PC1, PC2 and PC3) in the FP and HW were classified into categories by a density-based algorithm (see Methods, Supplementary Fig. S1-2) (*23*). The PCA predictions were compared to the ground truth of the histological analysis. Fig. 3a, b shows a color-coded comparison between the PCA clusters and the histological cluster, while Fig. 3c shows an example of the PCA prediction registered on a histological image. The comparison between PCA predictions and histological ground truth across the full dataset resulted in a sensitivity of 91% (60%), a specificity of 90% (100%) and an accuracy of 91% (82%) in the FP (HW) range (Fig. 3d) (Supplementary Fig. S3-6, see Methods).

**Figure 3.**
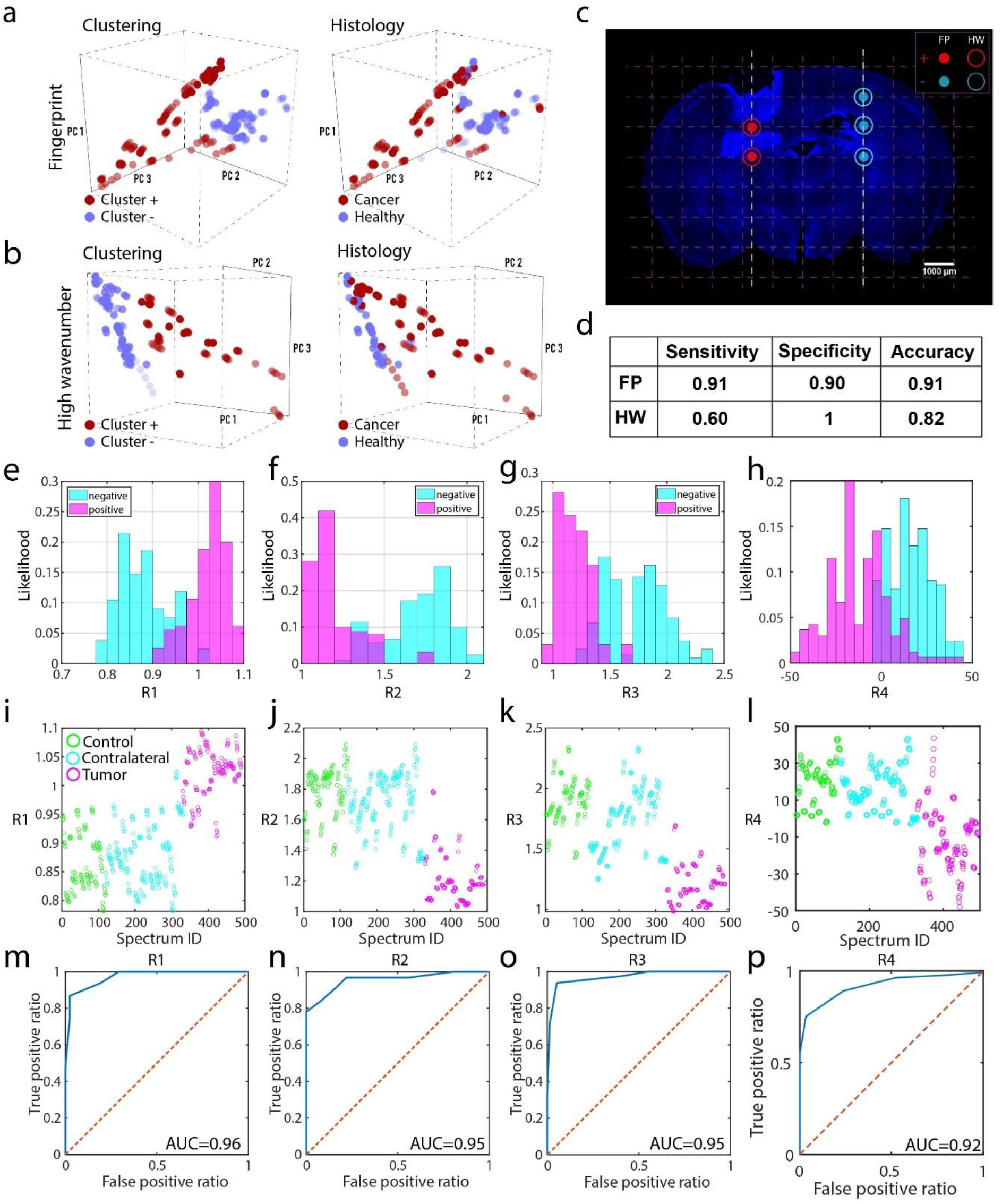
**a**, 3D plots of the Principal Components in the FP labelled following the PCA clusters (left) and the histology (right) **b**, as in a for the HW range **c)** Example of comparison between PCA clustering and histological imaging; dots and circles indicate the clustering in the FP and HW range respectively. **d**, Summary table of sensitivity, specificity and accuracy **e-h**, probability distribution of R1=S(1400)/S(1440), R2 =S(1440)/S(1800), R3 =S(1800)/S(2100), R4 =S′ (2847) +S′ (3100) where S(s) is the unprocessed Raman spectrum, s is the Raman shift and S′ = δS/δs; cyan, healthy tissue, magenta tumor **i-l** Scatter plots of R1, R2, R3, R4 for all measurements; green: control brain, cyan: healthy tissue in metastatic brain, magenta: tumor **m-p**, Receiver operator characteristics (ROC) for R1, R2, R3, R4 with Area Under the Curve (AUC).

We also devised a ratiometrics analysis based on salient spectral features B16/F10-BrM melanoma brain metastasis for rapid classification of the probed tissue (*24*) using the following figures of merit (FoMs), calculated on the raw Raman spectra *S(s)* (where s indicates the Raman shift) (Supplementary Fig. S7).

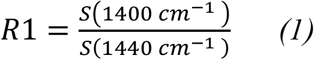

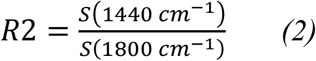

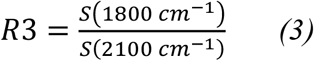

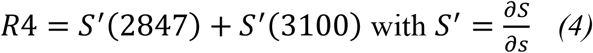

R1 reflects the intensity ratio between the peaks of melanin at 1400 cm^-1^ and lipids at 1440 cm^-1^. R2 mirrors the intensity ratio between the lipid band at 1440 cm^-1^ and the fluorescence tail at 1800 cm^-1^. R3 shows the presence of autofluorescence in the Raman silent region. R4 gives information on the strength of the onset and offset of CH bands respectively at 2847 cm^-1^ and 3100 cm^-1^ (Supplementary Fig. S8). We observed that the probability distributions of these FoMs’ values correlate with the histological classification (Fig. 3 e-h, see Methods). Also, the scores measured in metastatic regions deviate from the ones obtained in controls and healthy tissue in metastatic brains (Fig. 3 i-l). The suitability of these FoMs to discriminate between healthy tissue and metastasis is corroborated by having an Area Under the Curve (AUC) of the Receiver Operator Characteristic (ROC) comparable with Raman probes validated intaoperatively (*8*). We obtained an AUC of 0.96, 0.95, 0.95 and 0.92 for R1, R2, R3 and R4 respectively (Fig. 3 m-p).

Overall, these results demonstrate that minimally-invasive TF neural implants can detect tissue inhomogeneities as well as regions affected by metastatic cells in arbitrarily deep areas of the mouse. This can be done with sensitivity, specificity and accuracy, comparable with the state of the art of larger, non-implantable probes (*8, 21*). While this work is focused on metastasis from melanoma, our approach can be readily deployed to brain metastases from other primary tumors, such as lung cancer brain metastasis (Supplementary Fig. S9). At the same time, we suggest that the spatial resolution of the probe can be increased by fabricating optical collection windows on a metal layer coating the probe, reaching a resolution of hundreds of μm (Supplementary Fig. S10).

We believe that our method can provide experimental neuroscientists and cancer researchers with a promising complement to the existing set of technologies to interface with the deep brain. In particular, we envision that our approach might find relevant applications in neuro-oncology, where local therapies must be planned depending on the entity of the tumor (*25*). Owing to the peculiar capability of retrieving *label free* volumetric molecular information with low invasiveness, we also envision that our approach has significant potential to be deployed in organs other than the brain and in human subject, for example in detailed mapping of peritumoral regions for intraoperative margin assessment.

## Methods

### Tapered fiber implants

Tapered optical fibers were obtained from Optogenix s.r.l. and were fabricated from commercially available multimode optical fibers with NA=0.22, core/cladding size of 200/230 μm and low OH content. The tapers were produced by heat and pull as previously described (*26*). The fiber jacket was removed in acetone and the fiber stubs were connectorized with a metallic ferrule.

### Animals

All animal experiments were performed in accordance with a protocol approved by the Instituto Cajal CSIC, the CNIO, Instituto de Salud Carlos III and Comunidad de Madrid Institutional Animal Care and Use Committee. C57BL/6 mice 4–6 weeks of age were used. All protocols and procedures were performed according to the Spanish legislation (R.D. 1201/2005 and L.32/2007) and the European Communities Council Directive 2003 (2003/65/CE). Experiments were approved by the Ethics Committee of the Instituto Cajal and the Spanish Research Council.

### Brain sample for morphological Raman analysis

Adult C57BL/6 mice were obtained from the institutional animal facility. Mice were perfused with 4% paraformaldehyde in 0.1 M (pH 7.4) phosphate-buffered saline (PBS). Brains were post-fixed overnight, washed in PBS, and serially cut in 80 μm coronal sections (Microm HM450 sliding microtome). Sections containing the fiber tracks were identified with a stereomicroscope (S8APO, Leica) and mounted on glass slides in Mowiol (17% polyvinyl alcohol 4–88, 33% glycerin and 2% thimerosal in PBS) for posterior analysis.

### Brain samples with Melanoma Brain Metastasis

Murine B16/F10-BrM (brain metastatic) cells were all cultured in DMEM medium supplemented with 10% FBS, 100 IU.ml–1 penicillin/streptomycin, and 1 mg.ml–1 amphotericin B. Mice were injected intracranially with B16/F10-BrM (40000 cells in 2 μL) in the right hemisphere (1.5 mm lateral and 1 mm caudal from bregma, and to a depth of 1 mm) by using a gas-tight Hamilton syringe and a stereotaxic apparatus. After 11 days post injection, mice were anesthetized with CO2 and perfused with 4% paraformaldehyde (PFA). Whole brains were dissected and postfixed in the same fixative overnight at 4°C and then stored in phosphate-buffered saline (PBS).

### Raman Microscope

Raman spectra were acquired using a custom confocal Raman microscope. The TF was placed vertically in the focal plane of the microscope using a 3D micro manipulator. The fiber back facet was optically conjugated with both the output facet of a multimode fiber patch cord and with the entrance slits of a spectrometer. This was done using two optical paths separated by a dichroic mirror (785 Razor Edge dichroic, Semrock) placed in the infinity plane of a 4f system (Fig. 1c). In addition, the fiber back facet and the laser spot were imaged on a CCD camera to ensure optimal alignment. Laser light (785 nm, Lion Sacher laser) was filtered (785 MaxLine, Semrock) and relayed to the TF using a multi-mode fiber patch cord. The Raman signal emerging from the fiber was discriminated from the pump through the same dichroic and routed to a spectrometer (iHR320 Horiba Scientific Spectrometer with Horiba-Sincerity CCD cooled to *–*50 °C) after passing through an edge filter (785 Razor Edge ultrasteep, Semrock).

### Acquisition of Raman spectra

The mouse brain was placed on a 3D printed support whose corner, marked with a circle, was employed as a reference to identify the coordinates of the brain regions to be examined: tumor center, peritumoral regions and a healthy region. Moving the 3D support, we aligned the fiber tip with one of these areas and moved the brain upwards until the tip of the fiber touched the surface of the tissue. A lateral camera was used to ensure the brain surface was in contact with the tip of the taper. Prior to inserting the fiber into the brain, background spectra were recorded with the fiber in air. The TF was then inserted into the tissue in 1 mm increments, to a maximum depth of 3 mm from the surface. At each step, Raman spectra were collected through the same waveguide and guided onto the slit of spectrometer. The output power from the fiber was measured at 60-70 mW with an acquisition time of 30 seconds per spectra. The spectrometer slits were set to 200 μm and the grating at 600 lines per mm, while the spectrometer was controlled using Labspec Spectroscopy Software. Raman spectra in fig, 2d, e show averages of 5 spectra sequentially acquired in the same implant position in the FP and HW region after polynomial baseline subtraction.

### PCA Analysis

We analyzed Raman spectra using principal components analysis (PCA). PCA is a multivariate technique of analysis that can reduce the dimensionality of a dataset by finding the variables that contribute to its variance (*22*), i.e., the principal components (PCs). Each spectrum can be represented by some of its PCs without loss of information. Before the analysis, we pre-processed the data through polynomial base-line correction, min-max normalization and data resampling. Then, we processed the resulting spectra using custom-made algorithms demonstrated in previous reports (*22, 27, 28*). In brief, we determined the directions of maximum variance of the data as the eigenvectors of the covariance matrix of the spectra, and the principal components as the projections of the spectra onto this new subspace, following the protocol reported in reference (*28*). After the analysis, we found that the first three components (PC1, PC2 and PC3) account for more than the 99% of the information content of the original data, thus we used them to represent the spectra. Once the principal components were obtained, we plotted data in a PC1-PC2-PC3 diagram: in this graphic representation each point represents an individual measurement. This enables to find emerging patterns in the original data at a glance. Points were then clustered into groups using a density-based algorithm (*23*). The algorithm classifies elements into categories based on their similarity. Cluster centers were determined as those points with higher density than their neighbors and by a relatively large distance from points with higher densities. The steps of the clustering algorithms are recapitulated as follows. For each point *p* of the distribution: (a) its density *ρ* is determined as the number of points closer to *p* than the cut-off distance *δ*_*co*_. (*b*) *s* is determined as the subset of points with densities greater than the density of *p* - *ρ*(*s*)>*ρ*(*p*). (*c*) It is determined the point in *s* with minimum distance to *p*: this is the minimal distance of *p* from points with higher densities than *p* - *δ*_*min/ρ*_(*p*). (*d*) For each point of the originating distribution the density *ρ*(*p*) and *δ*_*min/ρ*_(*p*) are reported one against the other: this is the decision graph. (*e*) Cluster centers are spotted in the decision graph as the graph-outliers (points in the set with higher density than their neighbors and by a relatively large distance from points). (*f*) Points of the distribution are assigned to different clusters based on a smaller distance rule.

### Ratiometric Analysis

The values S(s) where averaged over 10 data points. The discrete derivative 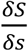 was calculated after a moving average of 20 data points. For all the FoMs, histograms were calculated by normalizing the sum of the frequency (entries per bin, n=160 total entries for the cancer group and m=210 for the healthy group) to 1 for both histologically defined classes. Table 1 reports average and standard deviation of the FoMs’ distribution. The null hypothesis that the two distributions of FoMs values that from negative and positive tissue belong to the same population has been rejected for all R1, R2, R3, R4 using a two-tailed T-test with unequal variances calculated as

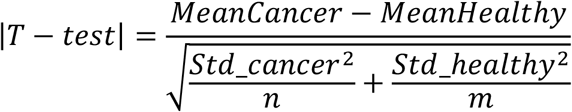

with significance α=0.01.

| FoM | Cancer | Healthy |  |
| --- | --- | --- | --- |
| | Mean ± Standard Deviation | | \|T-test\| |
| R1 | 1.02 ± 0.04 | 0.88 ± 0.05 | 28.94 |
| R2 | 1.20 ± 0.16 | 1.70 ± 0.19 | 27.56 |
| R3 | 1.19 ± 0.15 | 1.72 ± 0.026 | 24.65 |
| R4 | -12.15 ± 14.94 | 15.53 ± 11.57 | 18.77 |

The ROC curves were calculated by computing the ratios of false negative and false positive for an array of 10 values of the FoMs (e.g. R1), equally spaced between the minimum and maximum recorded values for each FoM.

### Histology and image registration

Slicing of the brain was done by using a sliding microtome (Thermo Fisher Scientific). 80 μm slices were blocked in 10% NGS, 2% BSA and 0.25% Triton X-100 in PBS for 1h at room temperature (RT) and then, nuclei were stained with bis-benzamide (1 mg/mL; Sigma-Aldrich) for 7 min at RT. Whole slices images were acquired with a Thunder Imaging System (Leica-Microsystems) equipped with a 10X NA 0.45 dry objective, with LED excitation, a DFC9000GTC camera and LAS X v3.5 software. The recording volumes were registered on the histological images using a frame of reference defined on the reference images acquired before every experiment (Supplementary Information). For each implant position, we considered three neighboring slices, each 80 μm thick. After retrieving the implantation axis and the depth of the TF tip at each measurement, the collection volume was determined considering a radius of r=250 μm around the taper axis surface and tip. The surrounding tissue was classified as positive (cancer invaded) if it belonged to the solid mass of the tumor and negative (healthy) if it did not show any sign of bis-banzamide staining.

*Classification* Sensitivity, specificity and accuracy were calculated as follows

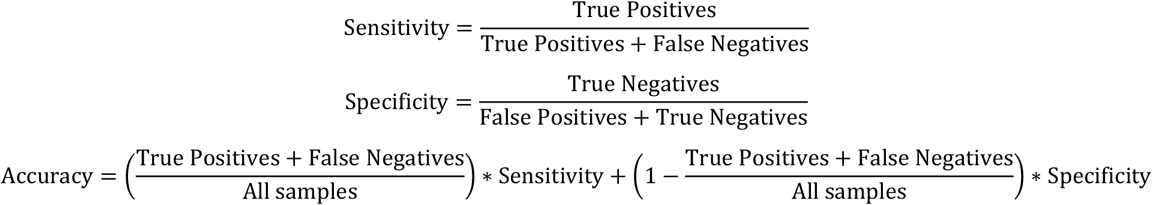

### Numerical simulations

The interaction volume of the TF probe was numerically simulated using a combination of Ray Tracing and Montecarlo techniques using Zemax software, following (*29*). Briefly, we modelled a TF stub as a straight portion of core-cladding multimode fiber and a conical section that was inserted in scattering medium. The straight portion of the fiber was modelled as two coaxial cylinders, 5 mm long, acting as core (radius R_1_=100 μm, refractive index n_1_=1.46) and cladding (radius R_1_=113 μm, refractive index n_2_=1.44). The taper cone was modelled using a single 2 mm long cone with base radius R_1_ and refractive index n_1_. The scattering medium was simulated using a Henyey-Greenstein model. To accommodate both excitation (785 nm) and collection wavelength (850-1050 nm) we used optical parameters reported for the first infrared window (*30*) using a scattering mean free path m=131 μm, scattering anysotropy g=0.92, transmission 0.9989 for emission at 785 nm and scattering mean free path m=158 μm, scattering anysotropy g=0.92, transmission 0.9989 for collection at 920 nm. 920 nm was chosen as the median wavelength between the FP and HW ranges limits. The collection map was calculated by scanning a point like emitter in the half-plane passing through the TF axis and measuring the power collected by the fiber with a bucket detector placed at the end of the straight portion. The emission map was calculated by injecting light in the full NA of the fiber from the straight end and by monitoring the emission power on a planar, pixellated detector containing the TF axis; the emission plot is normalized between 0 and 1. The photometry efficiencty for the TF was calculated as *Photometry* = *Collection x Emission* as previously described (*11, 16*). The white contour in the photometry plot (right) show the isoline of collection of 5% of the maximum collection value.

## Supplementary

### 1. PCA analysis

**Figure S1.**
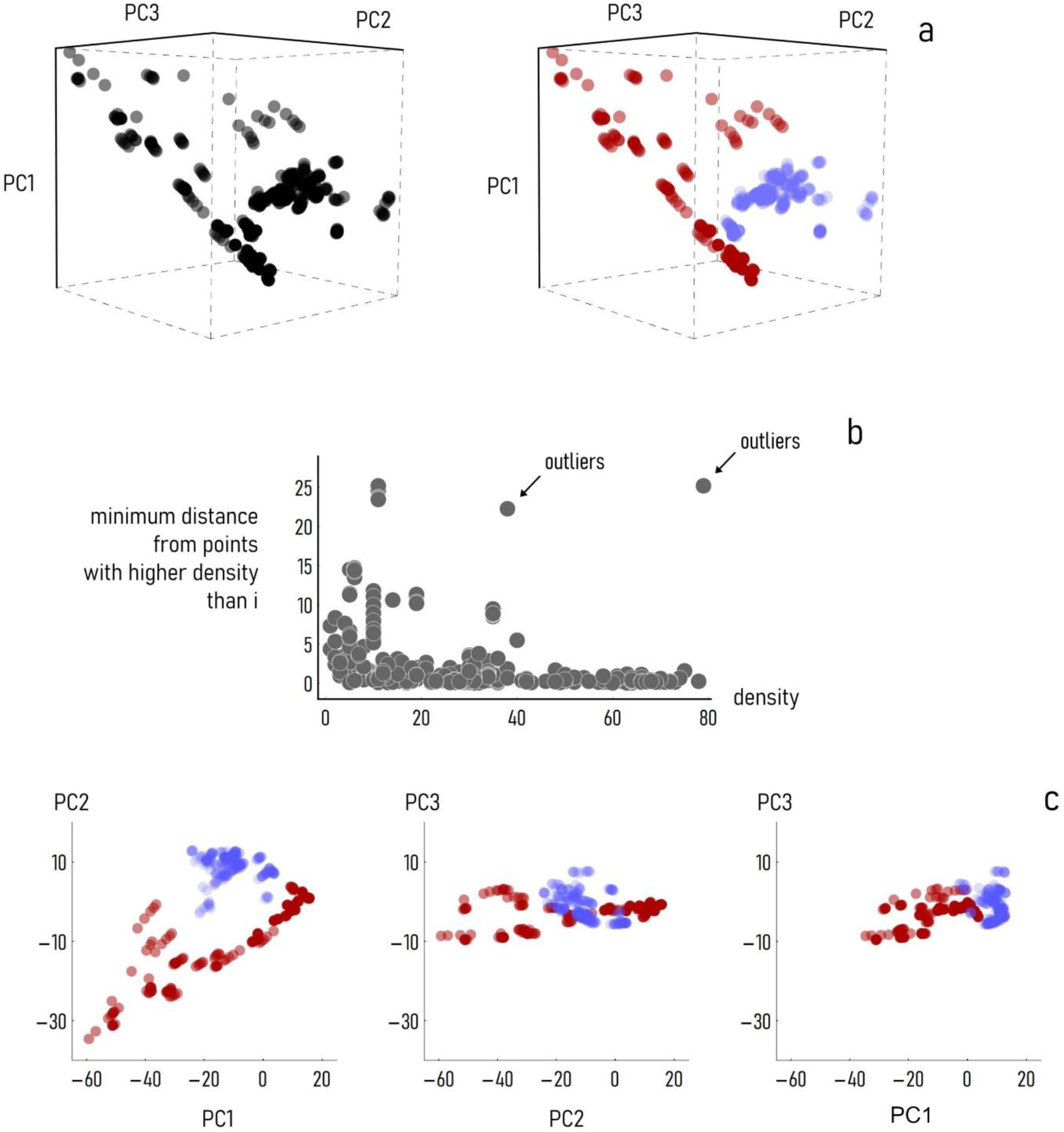
Example of PCA analysis in the LF range. a) PCA projections and clusters in 3 dimensions; b) Decision plot based on cluster density: the outliers identify the centers of the putative clusters; c) 2D orthogonal projections of PCA points.

**Figure S2.**
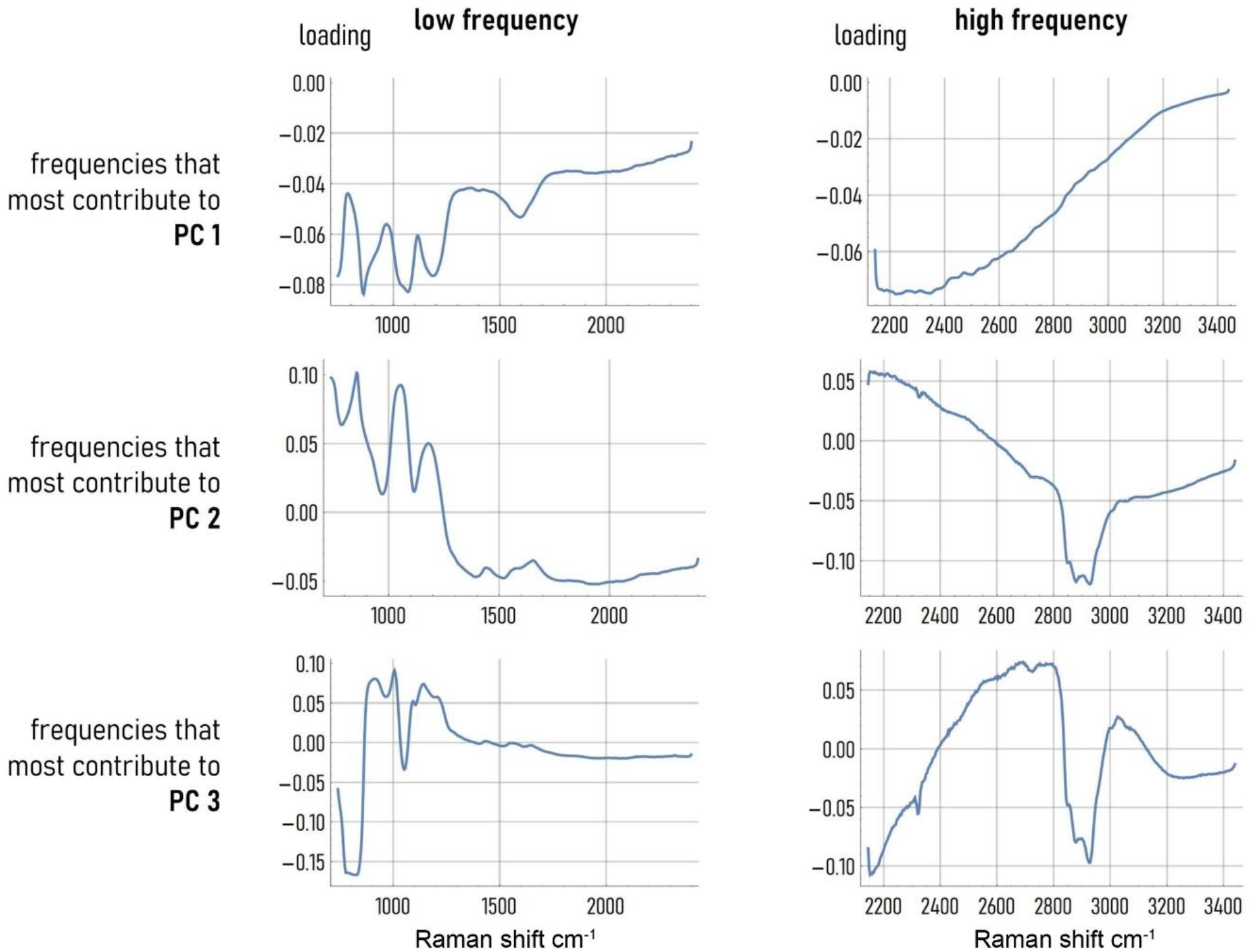
Detail of the three principal components extracted from the spectra in the fingerprint range (left colum) and high wavenumber range (right column)

### 2. Comparison of histological images and PCA predictions

This section illustrates the comparison between the PCA predictions based on the three dimensional clustering and the histology used as ground truth. Figures S3-6 show the histological images registered with the implant positions. For each measurement position in the metastatic brains, we classified the tissue as negative (cancer free) or positive (cancer invaded) taking into account the interaction volume of the fiber. This is shown graphically as a green or red shadow in the sampled region. For each of these positions, we overlayed the prediction of the PCA clusterings in the two frequency ranges (FP and HW). The clusters are color codified as Cluster +, cancer, magenta and Cluster - Healthy, cyan. The wavenumber range is codified as a filled circle for FP and ring for HW.

**Figure.**
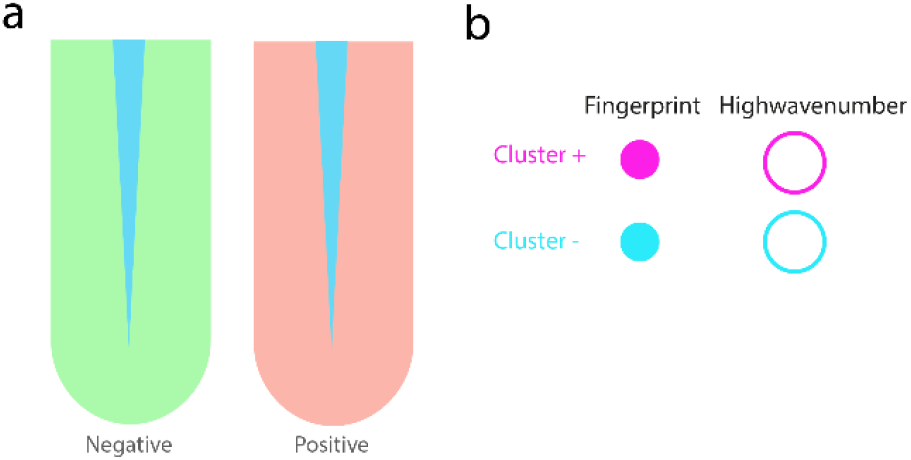
**a**, Legend of histological classification overlayed with histological images for healthy tissue (green) and tissue with metastasic cells (red) within the collection area of the probe **b**, Legend of PCA classification in cluster +, metastasis (magenta) and cluster -, healthy tissue (cyan) in the FP (dots) and HW (circles) ranges

**Figure S3.**
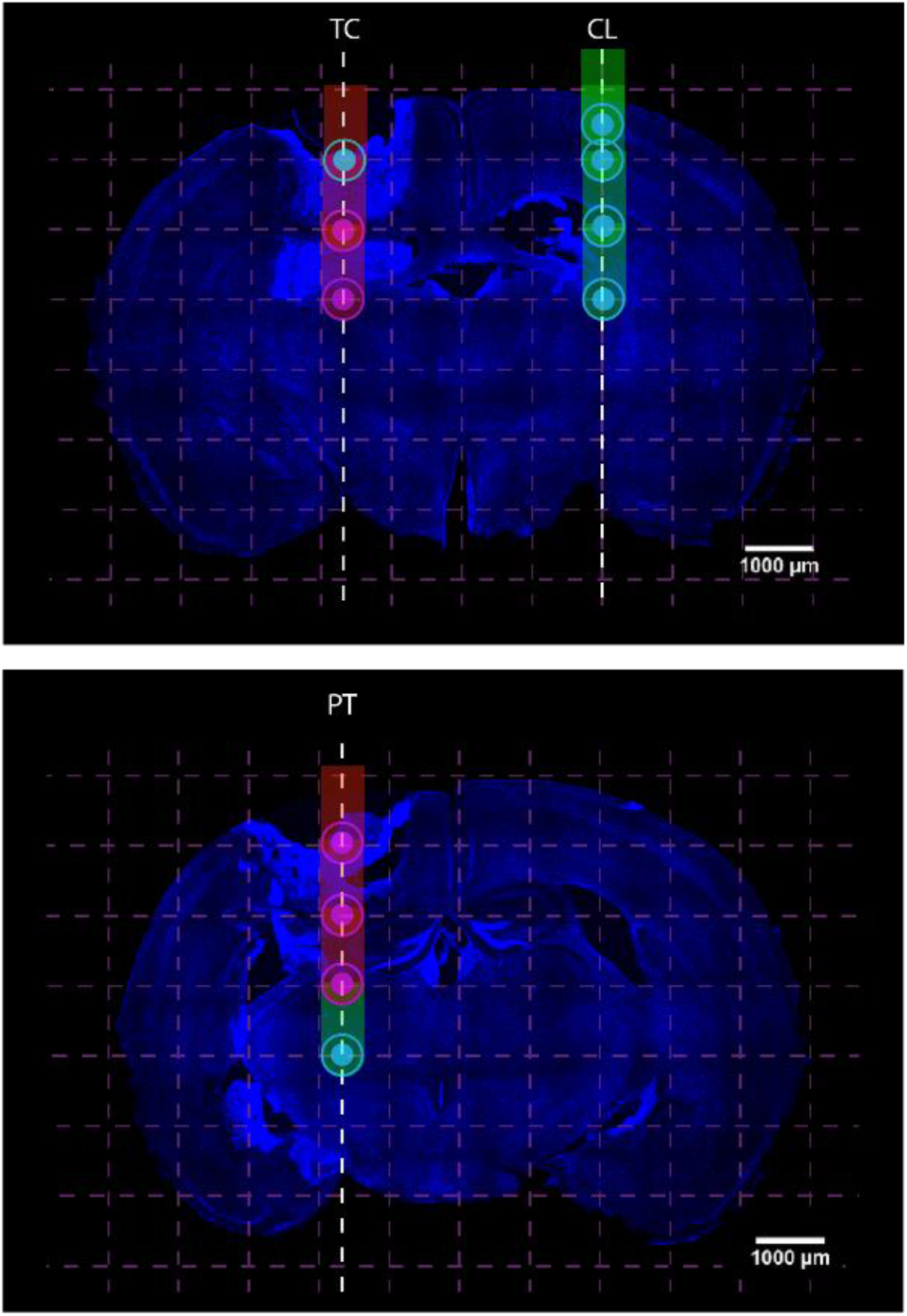
Histological images, classification and PCA clustering for Melanoma brain 1

**Figure S4.**
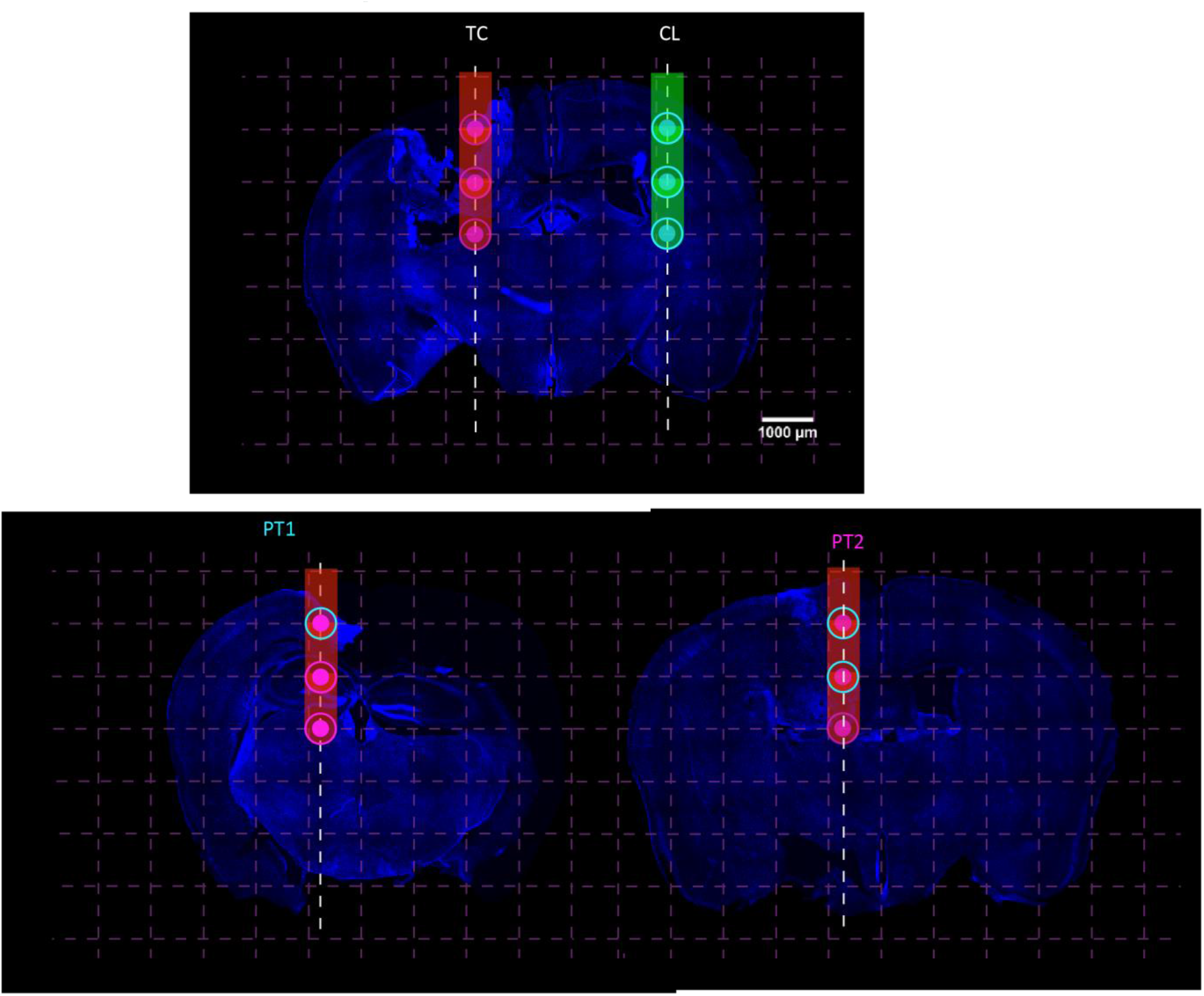
Histological images, classification and PCA clustering for Melanoma brain 2

**Figure S5.**
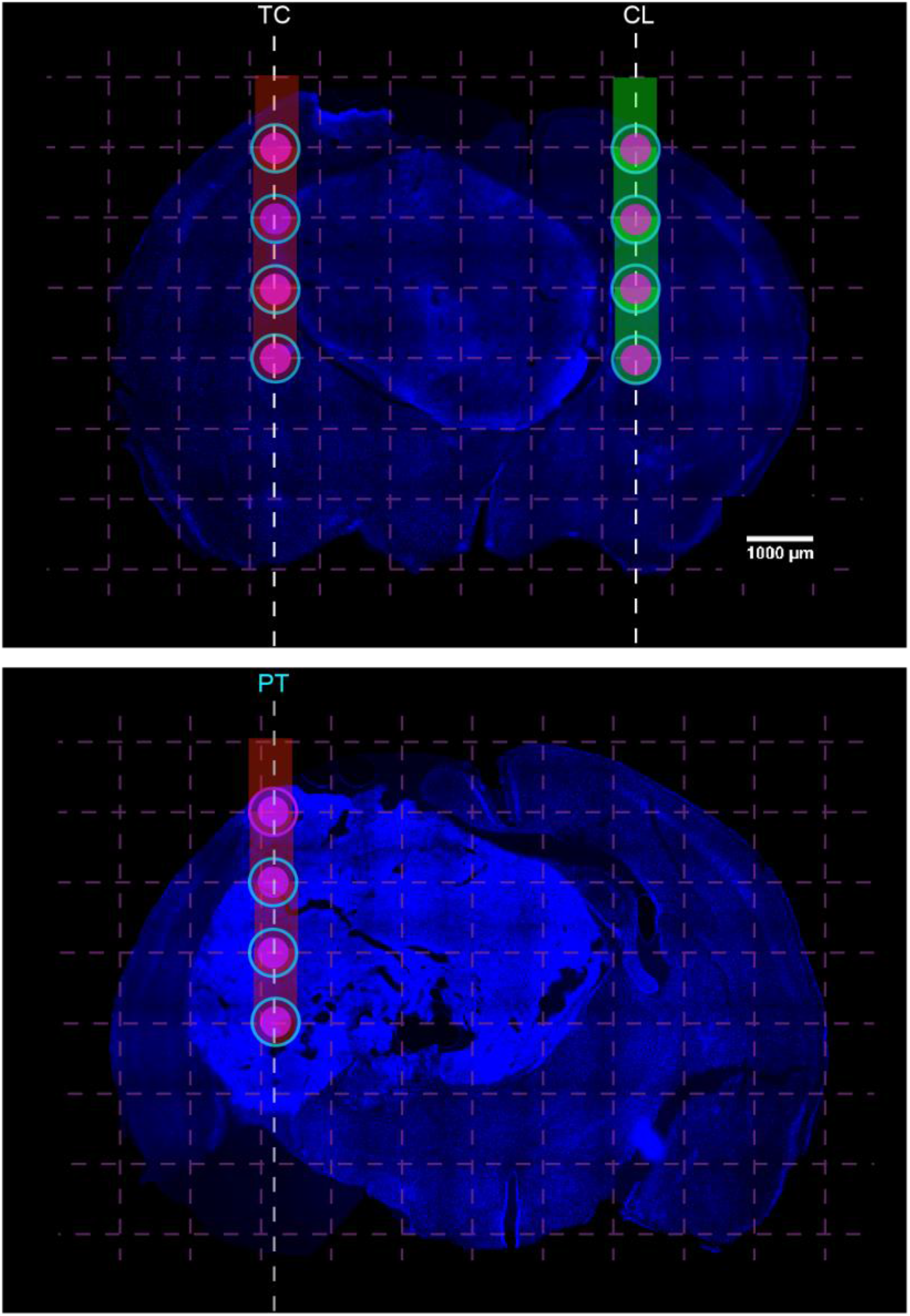
Histological images, classification and PCA clustering for Melanoma brain 3

**Figure S6.**
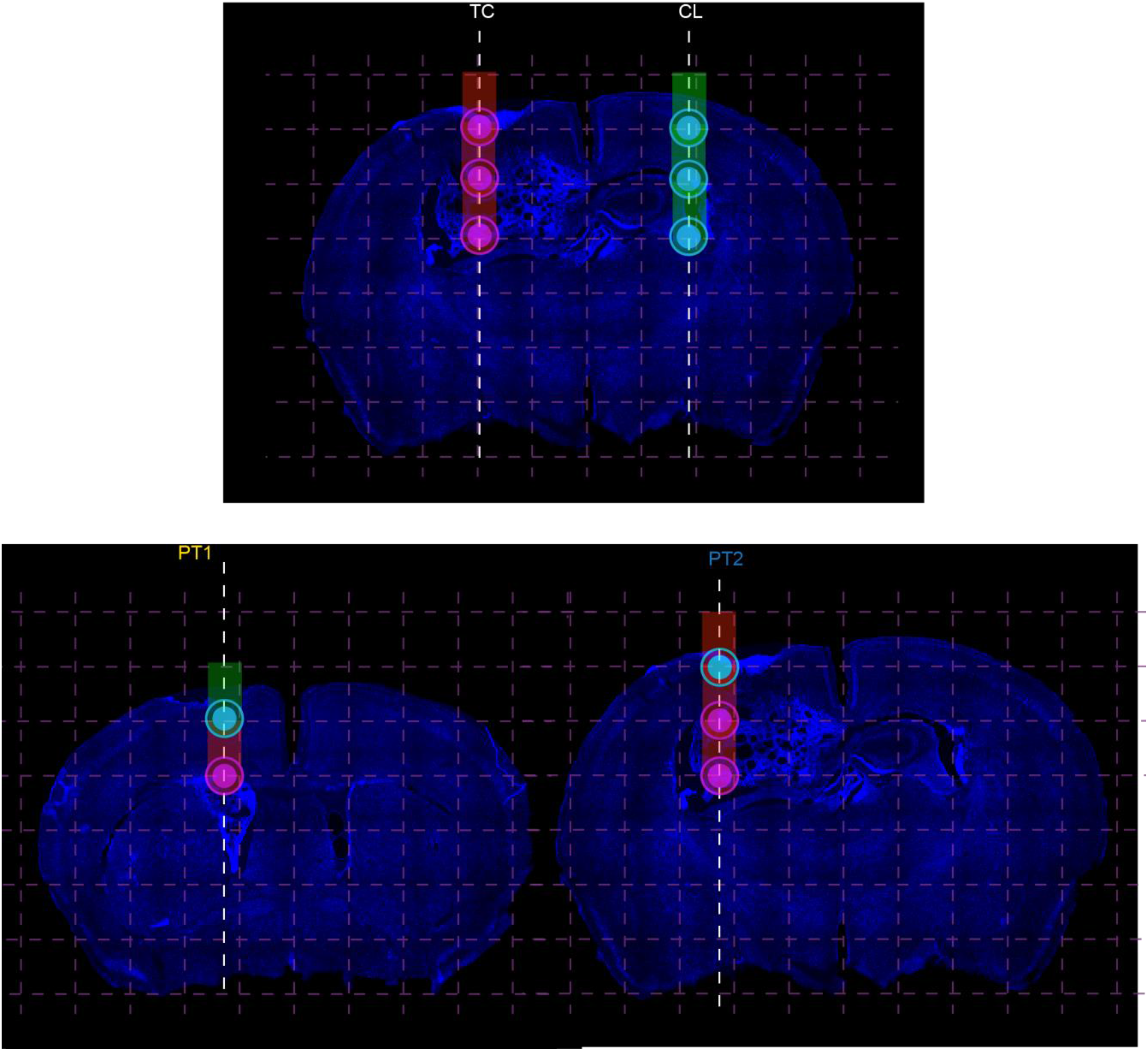
Histological images, classification and PCA clustering for Melanoma brain 4

**Figure S7.**
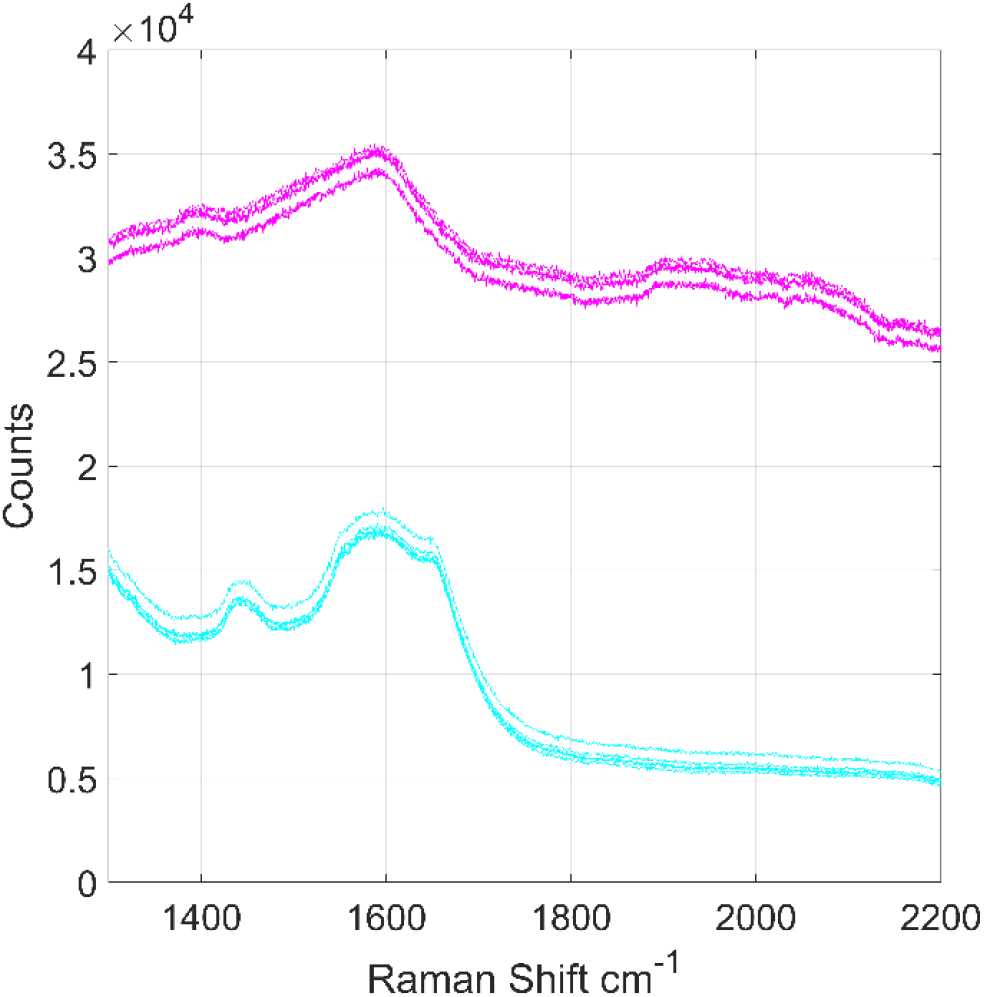
Example of the raw Raman spectra used for ratiometric analysis acquired in metastatic tissue (magenta) and in a contralateral position in healthy tissue (cyan).

**Figure S8.**
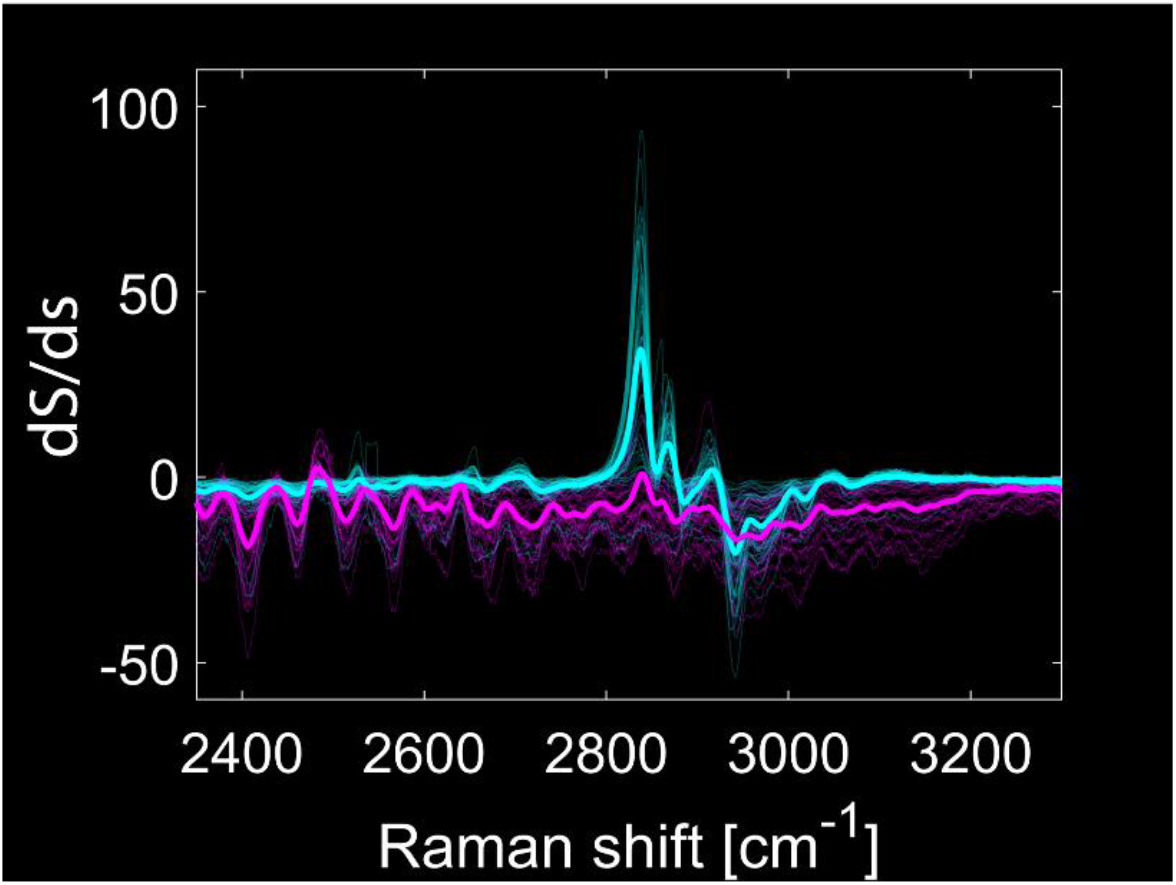
Derivative of Raman spectra for cancer free (cyan) and cancer invade (magenta) regions in the HW range. For display reason, the plot shows only 1 measurement out of 5 for every implant position.

**Figure S9.**
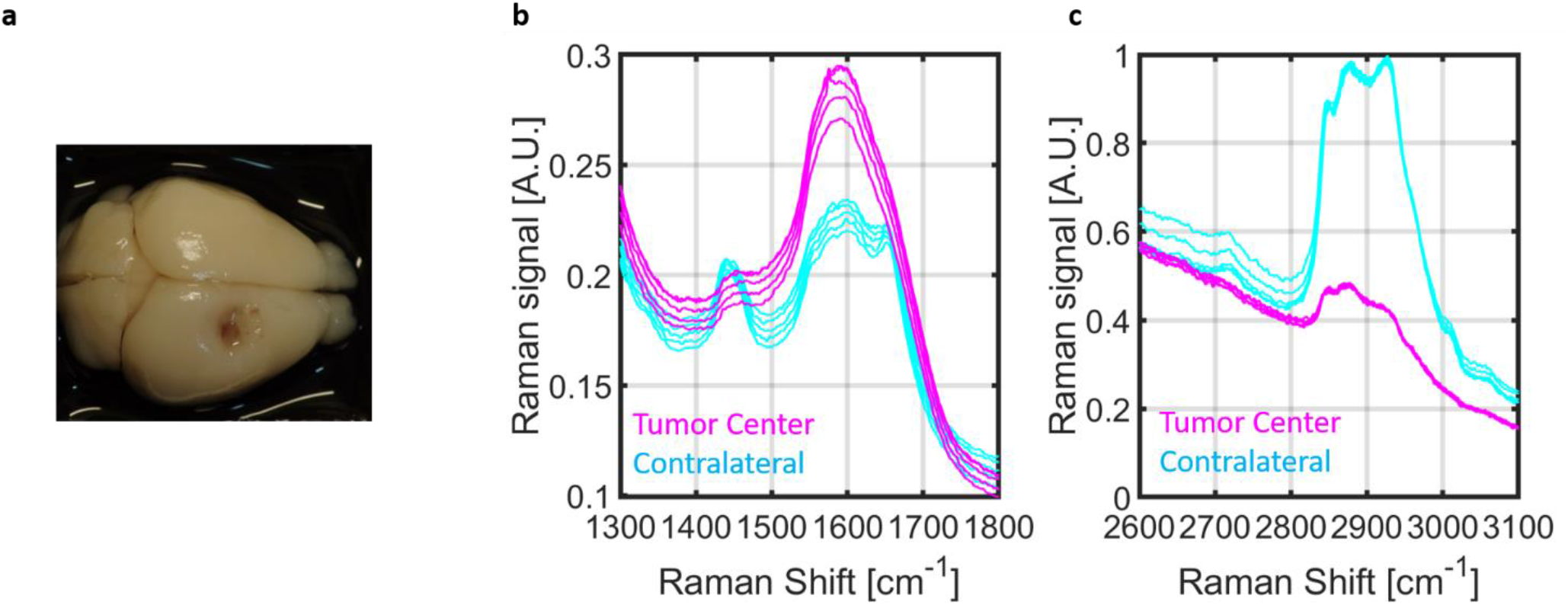
**a**, Image of a fixed mouse brain with an intracranially injected metastasis originating from cancer cell from lung primary tumor **b**, Raman spectra acquired in the center of the metastasis (magenta) and in the contralateral hemisphere (cyan) with the tip reaching -1 mm in the dorso ventral direction in the FP range. **c**, as in b for the HW range. Panels **b** and **c** show the spectral features that have been identified as reliable predictors of metastatic formation such as the suppression of lipid bands at 1440 cm^-1^ and 2847-3100 cm^-1^

**Figure S10.**
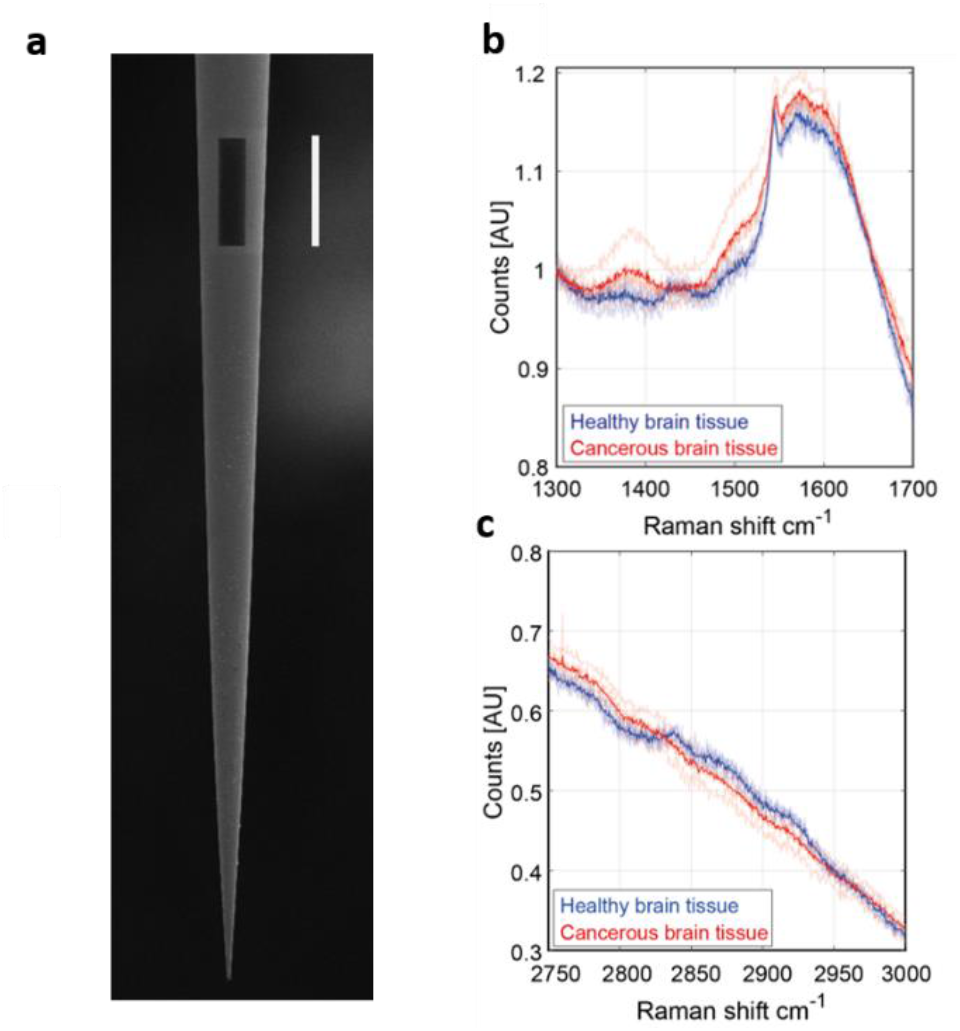
**a**, Scanning electron Microscopy (SEM) image of a tapered fiber covered by a thin gold layer (150 nm deposited by thermal evaporation with an optical window fabricated by locally removing the gold layer with low-current Focused Ion Beam milling (*31*); the scale bar is 100 μm. **b**, Raman spectra acquired in the center of the metastasis (red) and in the contralateral emisphere (blue) through the microstructured window the FP range; the thicker line show the average of 5 measurements **c**, as in b for the HW range. Panels **b** and **c** show the spectral features that have been identified as reliable predictors of metastatic formation such as the melanin peak at 1400 cm^-1^, the suppression of lipid bands at 1440 cm^-1^ and 2847-3100 cm^-1^.

